# Inferring the presence of aflatoxin-producing *Aspergillus flavus* strains using RNA sequencing and electronic probes as a transcriptomic screening tool

**DOI:** 10.1101/365254

**Authors:** Andres S. Espindola, William Schneider, Kitty F. Cardwell, Yisel Carrillo, Peter R. Hoyt, Stephen M. Marek, Hassan Melouk, Carla D. Garzon

## Abstract

E-probe Diagnostic for Nucleic acid Analysis (EDNA) is a bioinformatic tool originally developed to detect plant pathogens in metagenomic databases. However, enhancements made to EDNA increased its capacity to conduct hypothesis directed detection of specific gene targets present in transcriptomic databases. To target specific pathogenicity factors used by the pathogen to infect its host or other targets of interest, e-probes need to be developed for transcripts related to that function. In this study, EDNA transcriptomics (EDNAtran) was developed to detect the expression of genes related to aflatoxin production at the transcriptomic level. E-probes were designed from genes up-regulated during *A*. *flavus* aflatoxin production. EDNAtran detected gene transcripts related to aflatoxin production in a transcriptomic database from corn, where aflatoxin was produced. The results were significantly different from e-probes being used in the transcriptomic database where aflatoxin was not produced (atoxigenic AF36 strain and toxigenic AF70 in Potato Dextrose Broth).

## Introduction

Maize [1], peanuts [2], tree nuts, dried spices [3] and cottonseed [4] are crops that can be infected during the pre-harvest, post-harvest and/or storage period with *Aspergillus flavus* Link. This fungus produces polyketide secondary metabolites named aflatoxins. Among the four known aflatoxins (B_1_, B_2_, G_1_, G_2_), B_1_ has been of special interest to food biosecurity due to its toxicity and potent carcinogenic properties [5]. *A*. *flavus* is a ubiquitous saprophytic ascomycete fungus grouped in the *Aspergillus* section Flavi, species with aflatoxin-producing strains including *A*. *flavus*, *A*. *parasiticus* and *A*. *nomius* [6,7].

Aflatoxin is produced through the interaction of approximately 25 genes in a cluster cascade [8–10]. Regulatory genes for the cluster are *aflR* and *aflS* (*aflJ*), where *aflR* encodes for a transcriptional factor of the type Zn(II)_2_Cys_6_ which binds promoter regions of many aflatoxin genes [11–14]. In contrast, aflS (*aflJ*) regulates aflatoxin production through binding and activating aflR [15]. Some strains of *A*. *flavus* do not produce aflatoxin and these have been shown to have deletion mutations, identified by 32 separate PCR amplifications [16]. Callicott and Cotty [17] have begun to use cluster amplification patterns (CAPS) to evaluate *A*. *flavus* populations based on deletions genotypes.

Aflatoxin contamination in food is highly regulated in multiple countries, consequently increasing management costs and final product price [18–20]. In the United States alone, the maximum allowed concentration of aflatoxin in food for human consumption is 20 ppb, as dictated by the U.S. Food and Drug Administration (FDA). Appropriate and accurate aflatoxin testing is necessary to opportunely control *A*. *flavus* infected crops. Among the most used techniques for aflatoxin detection and quantification are thin layer chromatography (TLC), high-performance liquid chromatography (HPLC), enzyme-linked immunosorbent assay (ELISA) and fluorometry [18], however, there are limitations in all of these for rapid testing. Industry costs for testing crops for aflatoxins in the United States alone have ranged from $30 to $50 million per year at approximately at $10 to $20 per sample tested [20].

A promising management strategy for aflatoxin reduction is inundative biological control using atoxigenic strains of *A*. *flavus* (Aflaguard^®^ & AF36) [21–24]. Location or region-specific indigenous atoxigenic strains are selected to take advantage of inherent fitness in the environment and to avoid potential adaptation problems [25]. The indigenous isolated atoxigenic strains are multiplied *in vitro*, grown onto a carrier and inoculated on crop fields with the goal of competitive niche exclusion of the toxigenic strain, by the atoxigenic strains. The result is the almost complete elimination of aflatoxin contamination in the crop [21,26]. While the exact mechanism is not understood, a high prevalence of atoxigenic inoculum results in temporal and physical niche displacement of the toxigenic strains. Nevertheless, a single year release is not sufficient to render the field permanently atoxigenic. Thus a yearly application of the biological control agent is recommended for optimum efficacy [21,24,27].

Rapid assessment of determining changing profiles in atoxigenic to toxigenic population shifts have been lacking. Callicot and Cotty (2015) are attempting to create sensitive assays with cluster amplification patterns (CAPS), which involves some 32 known markers, however at this time; the CAP markers are primarily suggested to be a research technology for understanding dynamics of the atoxigenic populations. Absent that, current methods for assessing soil toxigenic to atoxigenic ratios of *A*. *flavus* are expensive, time, labor and skill intensive. Both, the evaluation of levels of soil *A*. *flavus* strain ratios and post-biocontrol screening could be facilitated by new methods. If there were a sensitive, simple assay for relative toxigenicity, inoculation of the field with additional atoxigenic strains could be recommended only when empirical tests showing higher relative toxigenicity reach an action threshold [28].

We propose that for practical application, the viability and presence of the toxigenic strains can be inferred by testing for upregulation of the gene cascade that leads to presence of aflatoxin. A fast and tentatively less expensive screening tool for toxigenic *A*. *flavus* strains might be sequencing the whole transcriptome (metatranscriptome) of the pathogen niche and determining the presence of toxigenic gene up-regulation. Previously, various approaches have been developed to use metagenomes in ecology studies and determine the microbial profile of natural ecosystems [29,30]. Yet, few have focused on the development of tools to detect microbes at the species/isolate level in agricultural ecosystems [31–33]. However, none of them has addressed the detection of gene activity and upregulation in agroecosystems. E-probe Diagnostic for Nucleic acid Analysis (EDNA) was designed to detect viruses, bacteria, fungi and oomycete plant pathogens by using species-specific markers named e-probes [31,32,34]. Here we modified EDNA to be used as a gene functional analysis and detection tool to infer the presence of aflatoxin. EDNA transcriptomics — a modification of the original EDNA’s bioinformatic pipeline — was designed to incorporate genome annotations on the e-probe design as well as on the detection pipelines. EDNAtran is a theoretical approach that is being tested for the first time with *A*. *flavus* and could be extended to detect metabolic functions associated to pathogenicity in other host-pathogen systems. Detecting metabolic functions that could potentially lead to plant disease is crucial to incorporate proper and timely management practices in agroecosystems.

## Materials and Methods

### Fungal isolates and culture methods

*A*. *flavus* strains were obtained as freeze-dried (AF36; ATCC 96045; atoxigenic) and frozen (AF70; ATCC MYA-384; toxigenic) cultures from ATCC (Manassas, VA). AF36 was reactivated by rehydration, adding 500 μL of sterilized distilled water inside the vial. Subsequently, 100 μL of the re-suspended AF36 was plated on Malt extract agar Blakeslee’s formula (MEAbl) and incubated at 31°C in darkness until mycelium was developed (72 hours), according to ATCC instructions. AF70 was thawed for 5 minutes, directly plated onto Malt extract agar, and incubated at 25 °C in darkness until mycelium was developed (72 hours), according to ATCC instructions. Agar plugs with actively growing mycelia were re-plated in MEAbl agar and incubated at their optimal temperatures in the dark until extensive conidial development (5 days) was observed. The cultures (AF36 and AF70) containing extensive conidia growth were used to inoculate ground corn and Potato Dextrose Broth (PDB).

Corn substrate was prepared using dried corn kernels (*Zea mays*). Kernels were weight (20g) and ground (using a coffee grinder Mr. Coffee Precision Coffee Grinder IDS77) until obtaining pieces with the approximate texture of coarse sand (0.5-1mm in diameter). The coarse grains were autoclaved (dry cycle) for 20 minutes in polycarbonate containers (Magenta GA-7, Plantmedia, US) and its humidity was adjusted to keep between 25 – 33% w/v (Modified from Woloshuk, Cavaletto, and Cleveland 1997).

Ground corn kernels and PDB media were inoculated with conidial suspensions obtained by washing *A*. *flavus* MEAbl plates with 2 mL of sterile distilled water. Conidia collected (2 mL) were then added to a single vial containing 4mL of distilled water for a final dilution of 3:1 v/v (Spore suspension was not quantified). Six mL of spore suspension was used to inoculate each replicate (20 g of ground corn and PDB). The ground grain was inoculated with the *A*. *flavus* suspension in polycarbonate containers and homogeneously mixed by rolling the containers to allow uniform distribution of the conidia. Similarly, 250 mL flasks containing 44mL of PDB were inoculated with 6 mL of *A*. *flavus* spore suspension. The containers and PDB plates were incubated at 31°C in the dark for 10 days.

### RNA extraction and sequencing

Ground corn kernels inoculated with the AF70 and AF36 strains produced extensive conidia, which were suspended by gently adding 10 mL of sterilized water to the magenta containers. The containers were shaken gently to homogenize the spores and then 1 mL of the spore suspension was obtained and added to a capped 2mL tube containing silica beads. The conidia cell walls were disrupted by shaking the 2mL tubes using a bead beater (2 cycles of 20 seconds). The lysate was then transferred (500 μL) to a column of the Qiagen RNeasy Plant Mini Kit for RNA extraction. On the other hand, for AF36 and AF70 growing on PDB, mycelia/spores were recovered by filtrating them using Whatman paper. 100 mg of mycelium/spores were weight and added to a column of the Qiagen RNeasy Plant Mini Kit to continue with the RNA extraction procedure. The RNA quality and integrity were assessed using a 2100 Agilent Bioanalyzer (Agilent Technologies) for 12 RNA extraction samples from AF36 and AF70. A sample (per strain) having RIN numbers higher than eight were selected for RNA sequencing. After quality control, RNA was sequenced using the Illumina HiSeq 2500 sequencer at the Core Facility of the University of Illinois at Urbana-Champaign, IL. The mRNA sequencing library was created with PolyA capture method per manufacturer’s protocol and the library was sequenced as single-end.

### Gene expression analysis

RNA sequencing reads from samples AF70-corn and AF70-PDB were mapped onto the *A*. *flavus* AF70 genome using STAR software [36] and bam binary files were transformed from sam files using SAMtools (http://samtools.sourceforge.net). Gene expression analysis was performed with DeSeq2 in R by comparing AF70 growing on two substrates (corn and PDB). The control was AF70 in PDB (non-conducive for aflatoxin production), and the treatment was AF70 in corn (conducive for aflatoxin production) [37]. Positive fold change (up-regulated) genes were selected using the log2 fold change metric obtained from the DeSeq2 analysis. Upregulated gene sequences having log2 fold changes greater than five were retrieved by an in-house Linux bash script and kept in a multi-fasta file for later e-probe design.

### E-probe design

The genomes from *A*. *flavus* AF70 (Accession: JZDT00000000.1) and NRRL3357 (Accession: AAIH00000000.2) [38] were obtained from Genbank. Sequences for the aflatoxin gene cluster of AF70 (AY510453) and AF36 (AY510455) were also retrieved from GenBank [39]. E-probes 80 nt long were generated using the e-probe pipeline for EDNAtran (Espindola *et al.*, 2015; Stobbe *et al.*, 2013). The aflatoxin gene cluster of AF70 was used as target sequence and the same gene cluster for AF36 was used as near neighbor sequence. E-probe specificity was verified by local alignment of each e-probes with the intended target genome (AF70) using an stringency of 100% identity and query coverage. Metadata information about the gene where it belongs was also retrieved by the EDNAtran e-probe design pipeline. E-probe annotation was utilized to select e-probes that belonged only to the up-regulated genes previously identified with DeSeq2.

### Rapid assessment of active aflatoxin metabolic pathway using EDNAtran

EDNAtran was utilized with default parameters (percent identity and query coverage of 100%) to assess four transcriptomic databases (Table 1) which included toxigenic (AF70) and atoxigenic (AF36) *A*. *flavus* strains growing in conducive (ground corn) and non-conducive (PDB) environment for aflatoxin production. E-probes designed in up-regulated genes of the aflatoxin gene cluster of AF70 was utilized during this analysis. Hit frequencies of raw reads with e-probes were recorded for each of the four treatments. Data on hit frequencies were analyzed for variance by ANOVA. For analysis of significant differences between hit frequencies, Tukey’s HSD test and T-test at P=0.05 were used.

**Table 1.** EDNA transcriptomics output table for the inference of aflatoxin in *A*. *flavus*.

| Culture | Total reads | ARL | LRL | MRL | Probe length | TNP | HQM |  |  |
| --- | --- | --- | --- | --- | --- | --- | --- | --- | --- |
|  |  |  |  |  |  |  | Ev | NoEv | HSGM |
| AF70<sup>a</sup> | 20657024 | 98.9 | 100 | 35 | 80 | 231 | 39 | 40 | 39 |
| AF36<sup>b</sup> | 22495368 | 99.0 | 100 | 35 | 80 | 231 | 2 | 2 | 2 |
| AF36<sup>a</sup> | 24134226 | 98.7 | 100 | 35 | 80 | 231 | 12 | 12 | 12 |
| AF70<sup>b</sup> | 24902500 | 98.9 | 100 | 35 | 80 | 231 | 231 | 231 | 231 |

ARL, average read length; LRL, largest read length; MRL, minimum read length; TNP, total number of probes; HQM, high-quality matches; Ev, matches include e-value in the scoring method; NoEv, matches do not include e-value in the scoring method; HSGM, high scoring general matches.
<sup>a</sup>*A. flavus* strain growing on PDB.
<sup>b</sup>*A. flavus* strain growing on ground corn.

## Results

### Assessing appropriate growing conditions for the production of aflatoxin

The isolates of *A*. *flavus* AF70 and AF36 showed different growth patterns and morphology on the different media (PDB and ground corn). Aflatoxin production is directly correlated with concentrations of free saccharides [40–43]; therefore AF70 and AF36 were grown in a toxin-inducing substrate (ground corn) and non-toxin inducing media (PDB). Free saccharides (glucose, sucrose, raffinose, etc) are localized in the germ of the corn grain when matured non-germinated seeds; however, in germinating corn, the endosperm is used as a source of saccharides [44]. Increased sclerotia production was observed in AF70, in contrast, AF36 produced conidia in all media. However, AF70 produced conidia 10 days post inoculation in corn.

### RNA sequencing and Gene expression analysis

RNA extracted from AF36 and AF70 strains grown on PDB and ground corn yielded from 20 to 24,9 million reads per sequencing run (Table 1). The sequenced reads then were mapped to the *A*. *flavus* AF70 strain genome to retrieve information about potential up-regulation and down-regulation of genes by using STAR [36] and DESeq2. In total, 44 genes were identified as up-regulated and 129 genes were down-regulated in the *A*. *flavus* genome (*Error! Reference source not found*. **Table**). Identified as part of the aflatoxin gene cluster, only 17 out of 44 upregulated genes. From two to six gene fold changes were plotted in a hierarchical clustering heat map as well as in a MAPlot (**Fig 1 Fig 2**).

**Fig 1. Mean Average Plot for RNA sequencing gene expression analysis. Red line shows zero change in gene expression**. Blue dashed lines show no change in gene expression and green dashed lines show a five-fold change in gene expression. Red dots are genes that have been either up-regulated or down-regulated in *A*. *flavus* AF70 infecting ground corn. Gray dots depict genes that have not had enough statistical evidence to be assigned a gene expression fold change.

**Fig 2. Hierarchical clustering map depicting *A*. *flavus* AF70 growing on PDA and ground corn.** Gene expression fold change is differentiated by a color palette ranging from red (most up-regulated genes have plus six-fold changes) to blue (most down-regulated genes have minus six-fold change). Genes are clustered based on their gene expression fold change to facilitate gene co-expression analysis.

### E-probe generation for afiatoxin detection

In total, 231 highly specific e-probes were generated to detect the production of aflatoxin specifically for AF70. AF36 genome-wide e-probes were not generated because there is not a genome sequence available yet for that specific strain.

### Detecting afiatoxin production using EDNAtran in A. fiavus

As expected, 231 e-probes had hits creating High-Quality Matches (HQMs) in AF70-corn transcriptome datasets; meanwhile, AF70-PDB had only 39 HQMs (**Fig 3**). AF36-corn had only two HQMs and AF36-PDB had 12 HQMs (Table 1). EDNAtran discriminated between the transcriptomic databases with abundant aflatoxin production and the transcriptomes from low-toxin production based on EDNA eukaryotic metrics [32] (Table 1). However, to infer the presence of aflatoxin we can only use frequencies of hits as an indirect measure. In this case, the number of times a read was mapped to an e-probe was recorded and counted without any limits. A dot plot of alignment length vs. percent identity with marginal hit frequencies facilitates visualizing hit frequencies. Specifically for *A*. *flavus* AF70 in corn, it was observable that the hit frequencies were very high — around 9,000 hits per e-probe — when the alignments are above 90% identity and the alignment length was approaching to the total length of the e-probe (**Fig 3A**). Conversely, for AF70 in PDB and AF36, the marginal plots show a low frequency of hits when alignment lengths and percent identities were above the threshold of 35nt and 90% respectively (**Fig 3B-3D**). Frequencies of hit values were square root converted and statistically analyzed with ANOVA to compare all the samples/treatments (**Fig 4**).

**Fig 3**. **EDNA transcriptomics hits distribution and frequencies for *A*. *flavus* aflatoxin detection**. (A and C) RNA sequencing of *A*. *flavus* AF70 and AF36 respectively growing on corn identified with 80-mer AF70 aflatoxin-specific e-probes. (B and D). RNA sequencing of *A*. *flavus* AF70 and AF36 respectively growing on PDB.

**Fig 4. Hit frequencies of AF70 e-probes in RNA sequencing databases of *A*. *flavus***. Atoxigenic strain of *A*. *flavus* AF36 growing on both PDB and Corn. Similarly, the toxigenic strain of *A*. *flavus* AF70 growing on PDB and Corn. Differences in gene expression levels are directly correlated to hit frequencies.

The ANOVA in the *A*. *flavus* experiment had a p-value lower than 0.05 which rejects the null hypotheses (all hit frequencies are equal); therefore, a post-hoc analysis was automatically performed using the Tukey HSD function in R. The post-hoc analysis and T-test for *A*. *flavus* showed that e-probes hitting on RNA sequencing databases obtained from *A*. *flavus* AF70 growing on ground corn were different from those of AF70 growing on PDB, and AF36 on corn and PDB (**Fig. 5** and Table 2). In conclusion, EDNAtran was able to find statistically significant differences between the transcriptomic data set of the highly toxigenic sample, from the non-toxigenic samples, using 231 e-probes generated in this study.

**Table 2.**
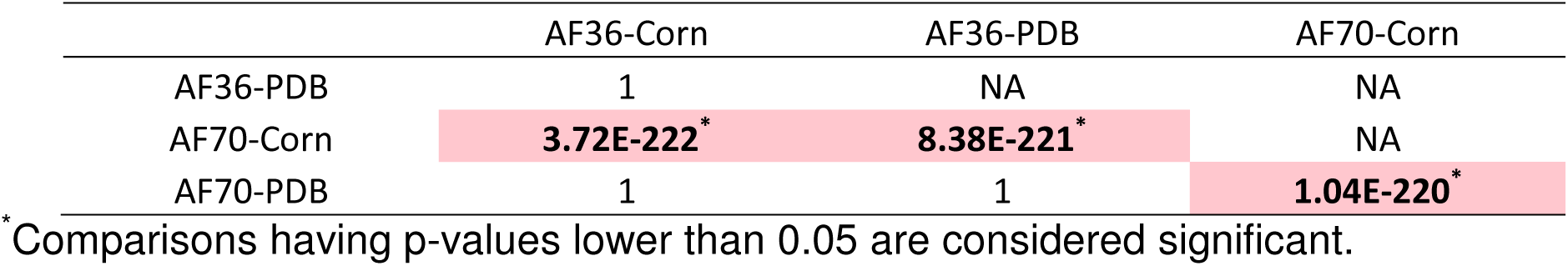
Pairwise T-test p-values comparing e-probe hit frequencies for *A*. *flavus* toxin detection analysis using the aflatoxin e-probes.

**Fig. 5**. Post-hoc analysis of ANOVA using Tukey HSD with 95% of confidence for the inference of aflatoxin transcriptional activation. Lines close to zero are interactions that had no difference in their hit frequency means while lines closer to 30 are interactions that had different hit frequency means.

## Discussion

EDNA has previously been proven to successfully detect a variety of plant pathogens from raw metagenomic databases [32,34,45]. DNA as the main source of identification has always been the gold standard for detecting organisms in a sample, although viability is not assessed. Therefore, the question about “dead or alive” is left undetermined unless the organism is isolated and cultured, or, transcriptome analysis is used as a complementary detection tool or, complementary molecular viability analysis is included [46].

Inspectors at international ports require a rapid detection method when decisions need to be done on site. EDNA has been considered a good candidate to be used as a diagnostic tool in ports of entry, due to its multiplexing capacity and rapidness. Yet, EDNA does not include an analysis of pathogen viability. If DNA-based detection (metagenomic analysis) is positive and viability needs to be addressed, the use of additional tests is not a viable approach for perishable or time-sensitive shipments. Using RNA sequencing and relative quantification of active genes is ideal to infer the viability of plant pathogens. The use of EDNA transcriptomics to infer the production of aflatoxin is a first attempt to introduce a novel strategy by using new sequencing technologies to identify viable plant pathogens. The use of e-probes that are designed on up-regulated genes incorporates an advantage to EDNA transcriptomics over other tools that use RNA sequencing to assess gene expression [37]. The advantage of EDNA transcriptomics over other methods of transcript frequency inference and calculation is that the time-consuming map against the reference genome is not necessary. Instead, we align the sample reads to the highly specific e-probes, which are designed for known up-regulated genes. Directing the analysis to genes that are known to be up-regulated reduces the analysis time tremendously since a mapping against a whole genome is no longer necessary. Where needed total nucleic acids (DNA and RNA) can be extracted from the sample of interest to perform both pathogen detection and gene activity.

Although most of the potential controlled inputs must be maintained constant, the sample matrix could contain fungal biomass, spores or sclerotia, depending on the organism and its life cycle stage. Yet, the source of relative quantitation become irrelevant because gene-expression analysis tools (including DeSeq.) are developed to analyze bulk populations — containing millions of cells —. Consequently, cell number differences between the treatment (Corn+AF70) and control (PDB+AF70) are small-uncontrolled inputs when equivalent sequencing depth has been achieved. Different cell counts between the treatment and the control in gene expression studies is therefore not a factor. We intend to use EDNA transcriptomics in metatranscriptomic analyses, where cell counts of organisms is difficult (or impossible i.e. unculturable-unknown organisms). Therefore, EDNAtran relies on a good quality sequencing data and equivalent sequencing depth to be able to differentiate between high and low-frequency hits.

The use of replicates in gene expression analyses using NGS are crucial, yet, this study used one replicate for all four RNA sequencing samples. In a real case scenario — where soil samples potentially containing *A*. *flavus* are collected — obtaining high-quality RNA libraries is more important than replicates. This study produced twelve RNA extraction samples from which four RNA extractions having the highest RNA integrity and quality (RIN≥8) where selected to be sequenced. Replication may be needed for statistical hypothesis-driven research, but it is not required for presence/absence queries or for the development of e-probes. Sequencing depth and sequencing quality equivalence are the most important metric for diagnosis.

Future studies need to include multiple blind samples to assess the usefulness of the new EDNAtran protocol to indicate the presence of aflatoxin-producing *A*. *flavus*. In this study, we have shown that in a known positive transcriptomic database, EDNAtran is capable of discriminating between production and no-production of aflatoxin. However, blind samples will provide a realistic assessment of the tool.

## Acknowledgments

This work was supported by the USDA-CSREES Plant Biosecurity Program, grant number 2010-85605-20542. Portions of the computing for this project was performed at the OSU High-Performance Computing Center at Oklahoma State University supported in part through the National Science Foundation grant OCI-1126330.

## Supporting Information

**S1 Table. Expression values (log2 fold Change) for AF70 *A*. *flavus* strain for two culture conditions (ground corn and PDB)**. For each Gene ID, expression levels are listed along with p-values.

